# APIPred Web 1.0: A Web Platform to Predict Potential Aptamer Sequences for Protein targets

**DOI:** 10.64898/2025.12.31.697194

**Authors:** Catherine Zhang, Juncheng He, Dhanush Gandavadi, Chau Nguyen Minh Hoang, Hyeongjun Cho, Minjun Son, Xing Wang, Abhisek Dwivedy, Saurabh Umrao

## Abstract

Aptamers are short single-stranded nucleic acids that bind protein targets with high specificity and are increasingly used in diagnostics and therapeutics, yet experimental discovery remains slow and variable in success. This creates a demand for computational systems that not only score candidate binders but also generate experimentally usable libraries under biologically meaningful constraints. Here, we present APIPred Web 1.0, a unified web platform that integrates constraint-aware aptamer library generation, machine learning- based aptamer- protein interaction prediction, and DNA secondary-structure analysis within a user-facing workflow. Users submit a target protein aminoacid sequence and define an aptamer template in a PREFIX - [VARIABLE] - SUFFIX format with real-time validation of key biological constraints (GC content and homopolymer limits). On the backend, sequences are converted into model- compatible features via optimized k-mer encodings (aptamer) and pseudo amino acid composition (PAAC) descriptors (protein), followed by inference with a trained XGBoost predictor. APIPred Web 1.0 improves the computational efficiency by applying precomputed protein features, vectorized batch processing (hundreds of sequences per batch), optimized XGBoost DMatrix inference, and a bounded heap that retains only the top 25 candidates during generation. The platform then computes minimum free energy (MFE) structures using ViennaRNA with parallel folding and returns ranked list of the top candidates with log-transformed interaction scores, complete sequences (variable region highlighted), dot-bracket structures, MFE values, and interactive 2D visualizations via persistent result links. In a demonstration study targeting CD64 protein, the platform produced 25 putative binders from a custom 40- nucleotide library and enabled selection of structurally diverse candidates for experimental testing. Flow cytometry showed specific binding to CD64-expressing THP-1 cells with minimal signal in Ramos control cells. Collectively, APIPred Web 1.0 offers a reproducible, structure-informed, and computationally efficient pipeline for rapid generation of aptamer candidates against target proteins for downstream experimental validation.

**Graphical Abstract:** 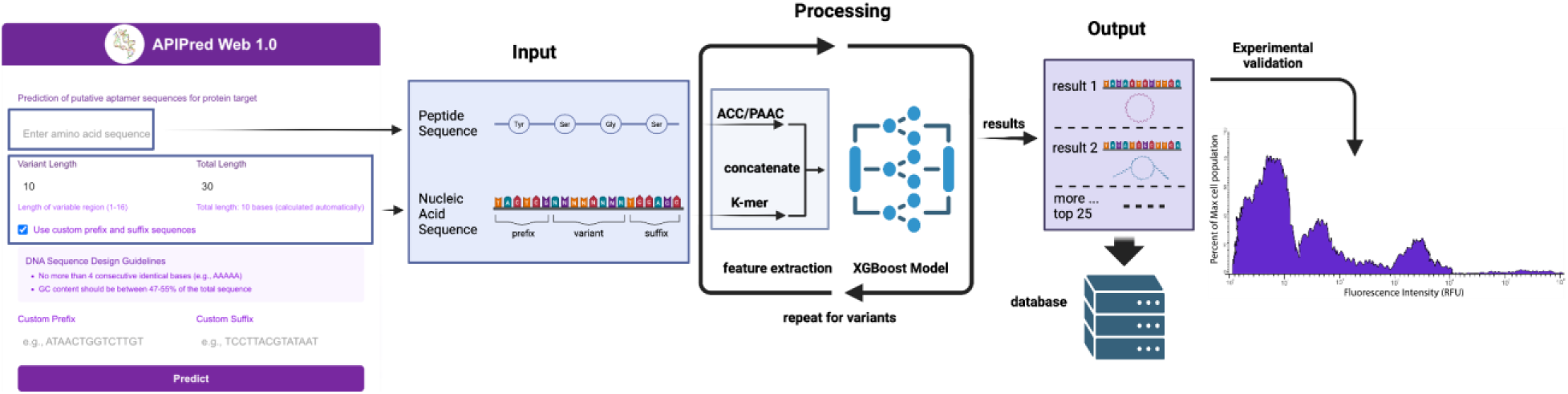

## Introduction

Aptamers are short single-stranded nucleic acids that recognize molecular targets through sequence-dependent folding into defined three-dimensional conformations. By forming shape- and chemistry-complementary interfaces, aptamers can achieve high specificity for protein targets and have therefore been explored across diagnostics, imaging, targeted delivery, and therapeutic inhibition. Compared with antibodies, aptamers offer practical advantages that are increasingly important for translation, for example, they can be synthesized chemically with high batch-to-batch consistency, readily modified with labels or functional groups, and typically display strong thermal stability and long shelf life. In addition, because aptamers are nucleic acids rather than proteins, they often exhibit reduced immunogenicity and can be engineered for controllable pharmacokinetics and reversible activity, depending on sequence design and chemical modifications. These attributes have contributed to strong commercial momentum in the space, with the aptamer technology market projected to expand from $3.63B (2023) to $26.23B (2034).^1^ Despite this promise, aptamer discovery is still dominated by SELEX (Systematic Evolution of Ligands by Exponential Enrichment), which requires iterative rounds of binding, partitioning, and amplification to enrich rare binders from large randomized libraries. SELEX performance is often limited by long development cycles (frequently weeks to months), sensitivity to experimental conditions (immobilization strategy, buffer composition, counter-selection stringency), and amplification/partitioning biases that can distort library composition over rounds. As a result, SELEX frequently demands weeks to months of iteration and optimization, and reported success rates can be low (often <30%), particularly for challenging targets or when high specificity under physiological conditions is required ^2^. These constraints motivate computational strategies that can reduce wet-lab burden by prioritizing candidates before synthesis and validation. To address these challenges, computational approaches for aptamer–protein interaction prediction have advanced rapidly. Early methods typically relied on feature representations (e.g., k-mer frequencies, nucleotide composition, physicochemical descriptors) coupled to classical classifiers or ensemble learners. More recently, the field has shifted toward deep learning models that learn hierarchical, task-optimized representations directly from sequences using convolutional networks to capture local motifs, recurrent networks to model order-dependent dependencies, and transformer encoders with self-attention to represent long-range contextual interactions. Current state-of-the-art systems like AptaTrans achieve ROC-AUC scores of 0.921 through transformer-based encoders with self-supervised pretraining. ^3^ In parallel, models including AptaNet ^4^ and hybrid CNN-LSTM systems like DeepAptamer ^5^ have further demonstrated the potential of deep learning-based architectures to improve prioritization of high-affinity aptamer candidates relative to experimental screening alone.

However, several practical and scientific gaps still limit routine adoption and downstream impact. First, existing web platforms such as PPAI can require complex feature extraction workflows, may be constrained in throughput for large candidate pools, and often provide limited interoperability between prediction, library design, structural analysis, and validation workflows ^6^. As a result, researchers often need to move between different programs for sequence generation, folding prediction, and experimental triage. Second, the field faces critical data quality issues, with 41% of publications reporting unexplained sequence alterations and insufficient standardization of evaluation metrics that complicate benchmarking and model generalization ^7^. Third, most current computational approaches focus on binary classification rather than quantitative affinity prediction, ^8^ while experimental decision-making often benefits from graded ranking and orthogonal parameters (e.g., thermodynamic plausibility and structural diversity) when selecting a small subset of candidates for experimental validation.

To address these limitations, we present APIPred Web 1.0, a comprehensive web platform that integrates advanced machine learning algorithms with optimized computational infrastructure to provide researchers with a unified, user-friendly interface for aptamer-protein interaction prediction, and secondary-structure thermodynamic analysis. Based on our published deep learning platform APIPred which combines feature engineering methods such as k-mer analysis, Pseudo Amino Acid Composition (PAAC), and structural modeling with backend optimizations that include parallel execution, vectorized batching, and more efficient data handling. Structural outputs are computed using the ViennaRNA Package 2.0 (PMID: 22115189) ¹⁰ to provide minimum free energy (MFE) estimates and dot-bracket structures alongside ranked interaction scores. Together, these improvements support higher accuracy predictions while keeping the system practical for routine research use by delivering biologically constrained candidate generation, scalable computation, and structure-informed interpretation in one reproducible workflow.

## Results

### 1. Generation of the Updated APIPred Algorithm

APIPred Web 1.0 is implemented as a two-tier architecture comprising a Python-based backend and a Next.js/TSX frontend. The backend is completely built with Python using several libraries to improve robustness and computational efficiency, which includes FastAPI, sqlite3, and others. An API (Application Programming Interface) is a defined way for different software systems to communicate, exchange data, and request services, while FastAPI is a modern Python web framework that makes it easy and very fast to build high-performance APIs. It integrates sequence processing, model inference, and secondary-structure analysis through a set of optimized computational modules. DNA folding and thermodynamic calculations are performed using the ViennaRNA package, and job metadata and results are stored in a local host server to support reproducibility and retrieval. To build the backend server, we first utilized the XGBoost model training Python script workflow from our published APIPred study and saved the trained model ^9^. We then refactored and optimized the feature extraction algorithm for high-throughput prediction. Specifically, the aptamer sequences are transformed using k-mer analysis, while protein sequences are represented using PAAC descriptors. The enhanced algorithm converts raw biological sequences into fixed-length numerical feature vectors that are compatible with the integrated predictive model, enabling comprehensive modeling of aptamer and protein sequence determinants relevant to binding The system implements several performance optimizations to make large-scale library screening practical in a web-server environment, and achieving significant computational speed. First, protein PAAC features are computed once per job and cached, eliminating redundant computations across candidate aptamer evaluations. Second, candidate evaluation is performed using vectorized batch processing (up to 500 sequences per batch) implemented with numpy arrays and optimized data structures, replacing inefficient sequence-by-sequence evaluation. We also vectorized constraint checking, which employs pre-compiled regular expressions and numpy operations for ultra-fast biological constraint validation, achieving order-of-magnitude improvements in filtering speed.

The prediction pipeline utilizes the XGBoost DMatrix API for optimized machine learning inference, providing 2-4x speedup over standard prediction methods through efficient memory layout and computation. During library evaluation, results are streamed into a bounded min-heap (priority queue) that retains only the top 25 candidates, providing O(log 25) insertion complexity. This prevents memory bloat and avoids expensive sorting operations on large candidate sets.For thermodynamic characterization, the backend performs parallel secondary-structure prediction using ViennaRNA with a ThreadPoolExecutor (typically four workers), yielding substantial speedups (3-4x) in folding throughput while maintaining robust execution

During sequence generation, APIPred Web 1.0 enforces biologically motivated constraints to promote experimentally tractable candidates. These constraints reduce the prevalence of sequences with extreme melting temperature or locally over-stabilized motifs that can drive misfolding and inconsistent DNA hybridization properties. The progress is calculated using weighted batch tracking with an exponential moving average to provide stable estimates of completion times. Upon job completion, the server returns the top 25 candidates ranked by predicted interaction probability, along with ViennaRNA-derived secondary structures and minimum free energy (MFE) values. All computed outputs and job parameters are stored in the SQLite database for later access and reproducibility.

### 2. APIPred Web Design - User Interface Components

The APIPred Web 1.0 frontend is implemented using the Next.js framework and provides an interactive interface for configuring prediction jobs and sequence libraries (Figure 1). The interface is designed to be accessible to users with diverse computational backgrounds while preserving the level of control required for experimental library specification. Users submit a target protein amino-acid sequence and define the aptamer library architecture that will be screened against the target.

**Figure 1.**
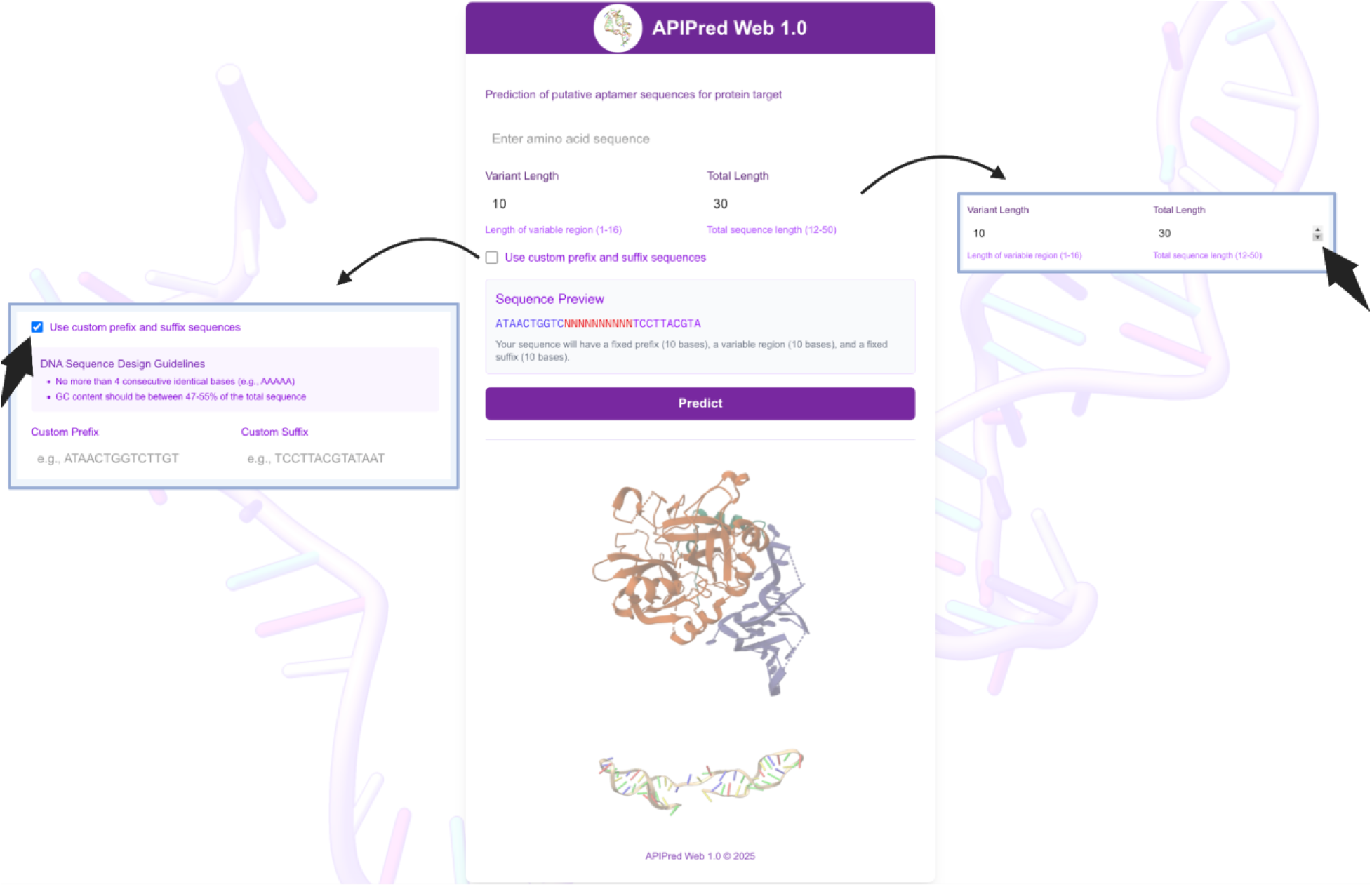
APIPred Web 1.0 interface for configuring the nucleic acid sequence pool. When an input field is selected, relevant interactive buttons become visible. Enabling the “Use custom prefix and suffix sequence” option displays an additional panel where users can specify custom prefix and suffix sequences.

As shown in Figure 1, the interface includes a required protein sequence input field as a required component, along with a dual-mode sequence design module. In default mode, the system automatically constructs prefix and suffix segments and allows users to specify a total sequence length (5-50 nt). In custom mode, users provide user-defined prefix and suffix sequences, which are validated in real time for allowed nucleotide characters and global design constraints. A variant-length selector (1-16 nt) controls the size of the combinatorial variable region. A live sequence preview renders the resulting PREFIX-[VARIABLE]-SUFFIX template with color-coded segments, enabling immediate visual verification of the intended library structure.

The platform incorporates comprehensive biological constraint validation with real-time GC content analysis ensuring the 47-55% requirement is met, consecutive base repeat detection preventing more than 4 identical bases, and consecutive G/C limitation enforcement preventing more than 4 G or C bases in sequence. Visual quality indicators provide immediate feedback with checkmark and error symbols to guide users toward valid sequence designs.

Upon job submission, the frontend communicates with the backend API to initiate an asynchronous prediction job and provides transparent execution monitoring. Job status is tracked via a polling-based progress system that reports percent completion and dynamically estimated time remaining using adaptive rate calculations based on observed processing speed, which also help users understand the computational intensity of their requests. Together, these UI components support a streamlined, end-to-end workflow from protein input and library design to computational screening and results retrieval.

### 3. Input Description and Requirements

The platform requires a continuous protein amino acid sequence as the primary text input provided as a plain string in standard single-letter notation. The sequence should be entered without spaces, special characters, or FASTA headers, and is treated as the target input for downstream feature extraction and prediction. For aptamers, users must define the length of the variable region (1-16 nts), with 10-12 nucleotides recommended for optimal balance of diversity and computational efficiency. **Figure 2** illustrates an example input to the web interface, showing how an amino acid sequence is entered and how the corresponding nucleic acid sequence composed of a fixed prefix, a combinatorial variable region, and a fixed suffix (PREFIX-[VARIABLE]-SUFFIX). The total length configuration operates in two modes: default mode, which allows users to specify a total sequence length of 5-50 total bases with automatic prefix and suffix generation, while custom mode enables user-defined prefix and suffix sequences that must contain only A, T, G, C bases and meet all biological constraints.

**Figure 2.**
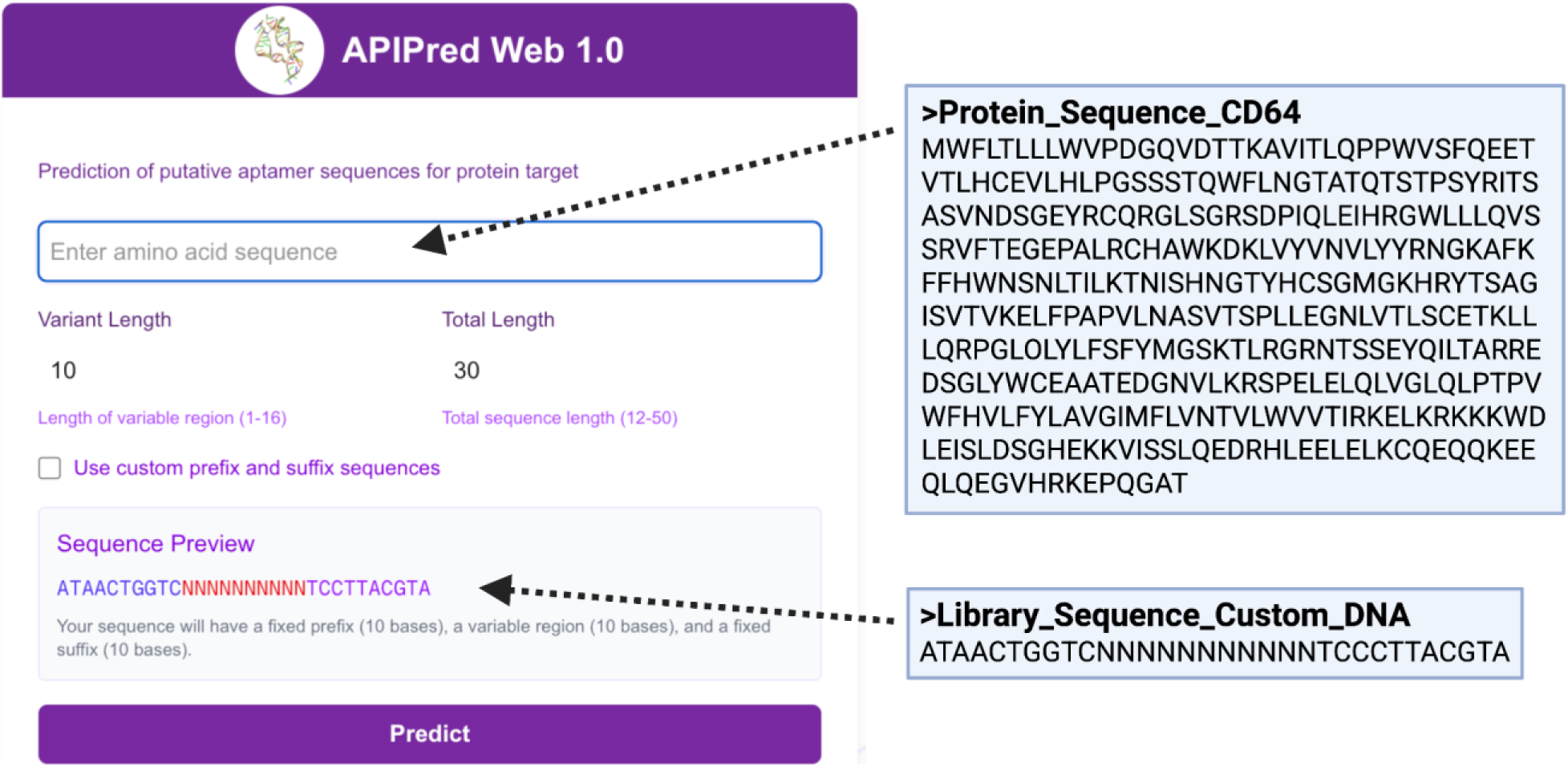
Illustration of an example protein input and aptamer sequence pool structure used by APIPred Web 1.0. The protein sequence input is provided as a plain single letter amino acids string. The aptamer library is specified using standard single-letter DNA bases (A, T, C, G) and generated in a PREFIX-[VARIABLE]-SUFFIX format for downstream screening.

The default library parameters use fixed flanking sequences (prefix: ATAACTGGTCTTGTTACAGGTCTG; suffix: TCCTTACGTATAATACTACCGAAC). Custom sequences must satisfy two critical biological constraints: GC content maintained between 47-55% across the entire sequence and no more than 4 consecutive identical nucleotides. To ensure interactive performance and predictable runtime, the platform limits each job to a maximum of 50 nucleotides total sequence length and a maximum of 1,000,000 candidate sequences per request, with a default library size of 65,536 sequences for routine analyses.

### 4. Output Interpretation - Sequence, Score, and Structure Analysis

APIPred Web 1.0 returns a ranked list of candidate aptamers together with quantitative prediction and structural annotations. The primary output provides interaction prediction scores reported as log scores, which represent the natural logarithm of model-predicted interaction probabilities. ln (p). This log transformation makes high-ranking probability predictions more readable, converting values like 0.99989 to more interpretable log values such as −0.0000478. In this scale, higher (less negative) log scores indicate stronger predicted protein-aptamer interactions, and candidates are ranked in descending order of log scores to present the best probable aptamer candidates first.

In parallel, APIPred Web 1.0 reports thermodynamic and structural properties derived from DNA secondary-structure prediction. For each top-ranked candidate, the platform computes the MFE and corresponding secondary structure using ViennaRNA dynamic programming algorithms parameterized by the Turner 2004 energy model.. More negative MFE values indicate greater thermodynamic stability of the DNA secondary structure, with typical values ranging from 0 to −50 kcal/mol for sequences of 20-50 nucleotides. The structure representation uses dot-bracket notation to show base pairing patterns. It is important to note that MFE and interaction probability scores are independent measures. The interaction score reflects the learned likelihood of target binding, whereas MFE reflects intrinsic folding stability. Accordingly, the high interaction probability does not necessarily correlate with a strongly stabilized (more negative)MFE, as binding affinity depends on protein-specific interactions rather than general structural stability.

For each candidate, the results table displays the full aptamer sequence in PREFIX-[VARIABLE]-SUFFIX format, with the variable region highlighted to facilitate comparison across candidates. The platform additionally provides interactive 2D structure visualization of predicted folds using the Forna library and D3.js, enabling users to inspect base-pairing geometry with color-coded nucleotides and interactive features like hover effects and structure manipulation capabilities. Figure 3 illustrates how these outputs are presented in the results interface.

**Figure 3.**
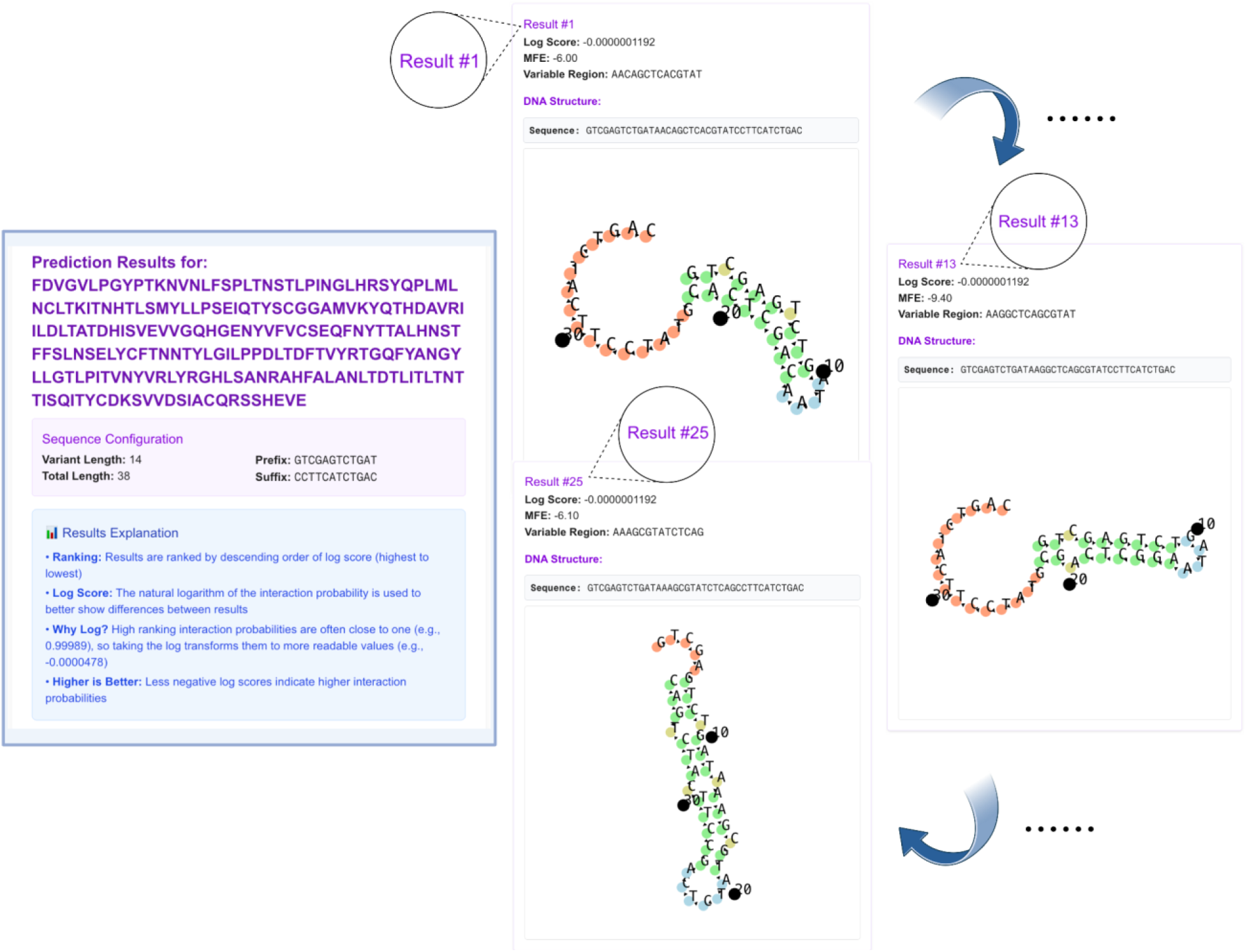
Example output page generated from a completed job. **Below** the configuration summary, the interface displays the top 25 predicted sequences of all possible outcomes. Each result panel reports the ranking, log score, MFE, the complete nucleic acid sequence with its variable region, and an interactive rendering of the predicted 2D secondary structure.

To support efficient use in exploratory settings, APIPred Web 1.0 includes a user-controlled termination feature that allows researchers to halt lengthy computations while still retrieving partial results. Besides the basic interface components and interactions with the backend server API, we integrated several Forna containers in combination with D3.js for visualizing the 2D structure of the best-match aptamer sequences computed from the frontend. For users’ convenience, we provide a module for previewing the structure of the DNA sequence that will be generated, with clear guides on the length and structure of input displayed on the interface. Warnings appear when the input is not legitimate, and the input is validated in real-time. When the estimated time is too long, a notification appears to inform the user. We provide clear guidance on accessing results through unique and bookmarkable URLs that remain activefor 30 days following job completion.

### 5. Example Aptamer Prediction and Experimental Validation

To evaluate the accuracy of the prediction framework, aptamers targeting CD64 were generated. CD64, also known as Fc gamma receptor I (FcγRI), binds monomeric immunoglobulin G (IgG) and is ubiquitously expressed on monocytes and macrophages, where it plays critical roles in pathogen recognition and clearance, as well as in the targeting of cancer and pathogen-infected cells. APIPredWeb was provided with the amino acid sequence of CD64 along with a custom-designed 40-base aptamer library (**Figure 4a**). The library comprised a central region of 10 variable nucleotides flanked by a default 15-base prefix and 15-base suffix sequences. These sequences were designed to maintain an approximate GC content of 50% and melting temperatures ranging from 55–60 °C. Within 72 h, APIPred Web predicted 25 putative CD64-binding aptamer sequences. These candidates were subsequently analyzed for predicted secondary structures, and 10 aptamers with distinct and nonredundant structural features were selected for experimental validation (**Figure 4b**). The secondary structures of the selected aptamers are shown in **Figure 4c**.

**Figure 4.**
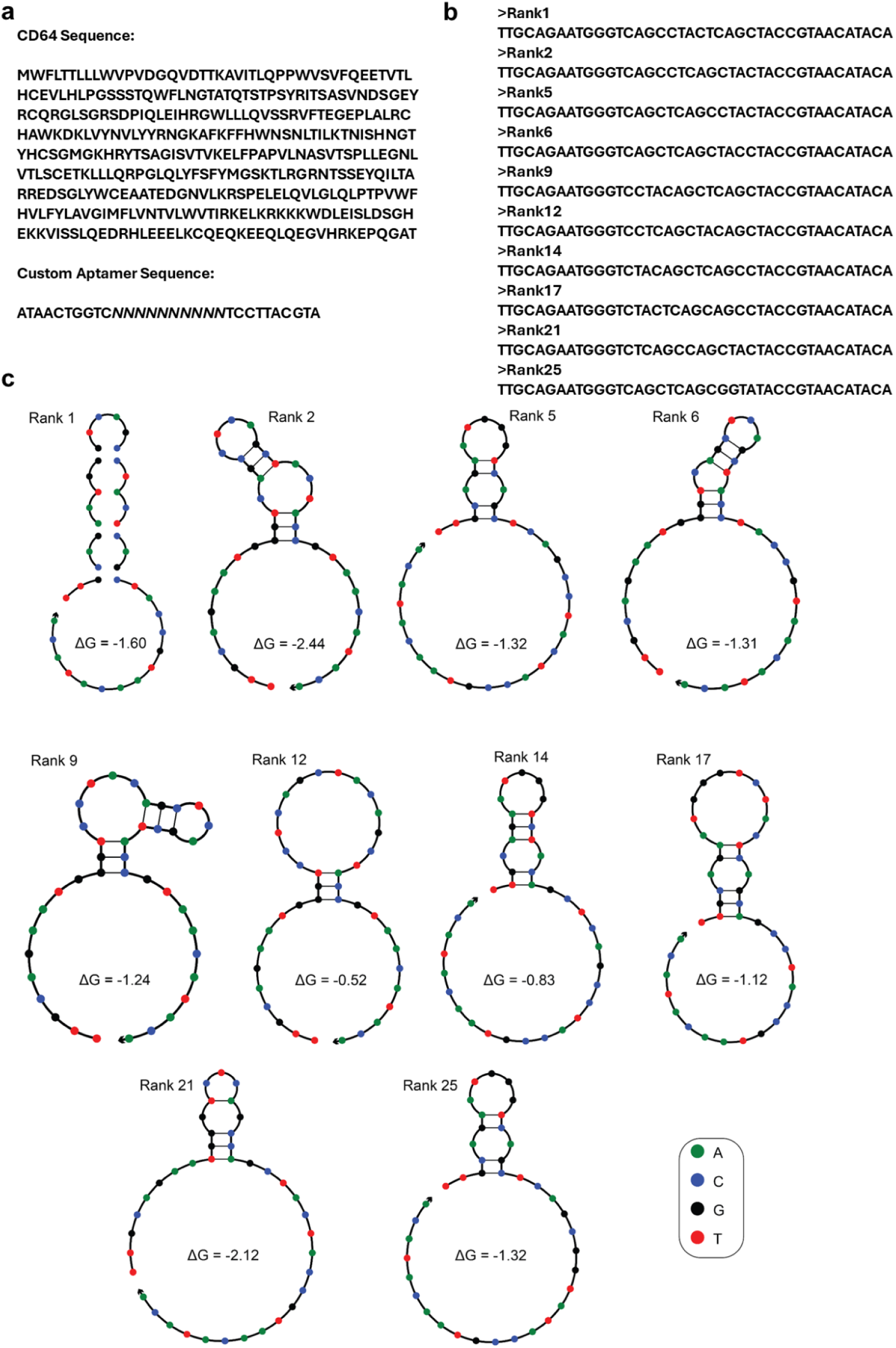
Prediction of aptamers for CD64. **(a)** Input protein sequences for CD64 (PDB/Accession no.?) and 40 bases long aptamer library with randomly generated prefix and suffix sequences with 10 variable bases. **(b)** Sequences of 10 aptamer chosen from APIPred output with unique secondary structures. **(c)** Secondary structures of the 10 chosen aptamer showing unique structural elements and folds.Gibbs free energy (ΔG) values are reported in kcal/mol.

The selected aptamers were obtained from a commercial vendor with a 5-FAM fluorophore conjugated to the 3′ end. THP-1 (monocytic) cells were used to evaluate aptamer binding, whereas Ramos (B lymphocyte) cells, which exhibit minimal CD64 expression, served as negative controls. Flow cytometry was employed to quantitatively assess aptamer binding to both cell lines at an aptamer concentration of 10 µM. All predicted aptamers demonstrated detectable binding to THP-1 cells **(Figure 5a - i)**, with Apt1, Apta2, Apta9, Apta12, and Apta21 exhibiting particularly strong binding, yielding fluorescence intensities on the order of 10⁵ following background correction. In contrast, Ramos cells displayed negligible fluorescence signals (on the order of 10⁵) after background subtraction, indicating minimal nonspecific binding (**Figure 5j**). Although APIPredWeb-based ranking generally correlated with binding performance, further validation by end users may be warranted, as lower-ranked aptamers (e.g., Apta21 for CD64) also exhibited strong and specific binding. Collectively, these results suggest that APIPredWeb effectively predicts aptamers with high binding affinity and target specificity.

**Figure 5.**
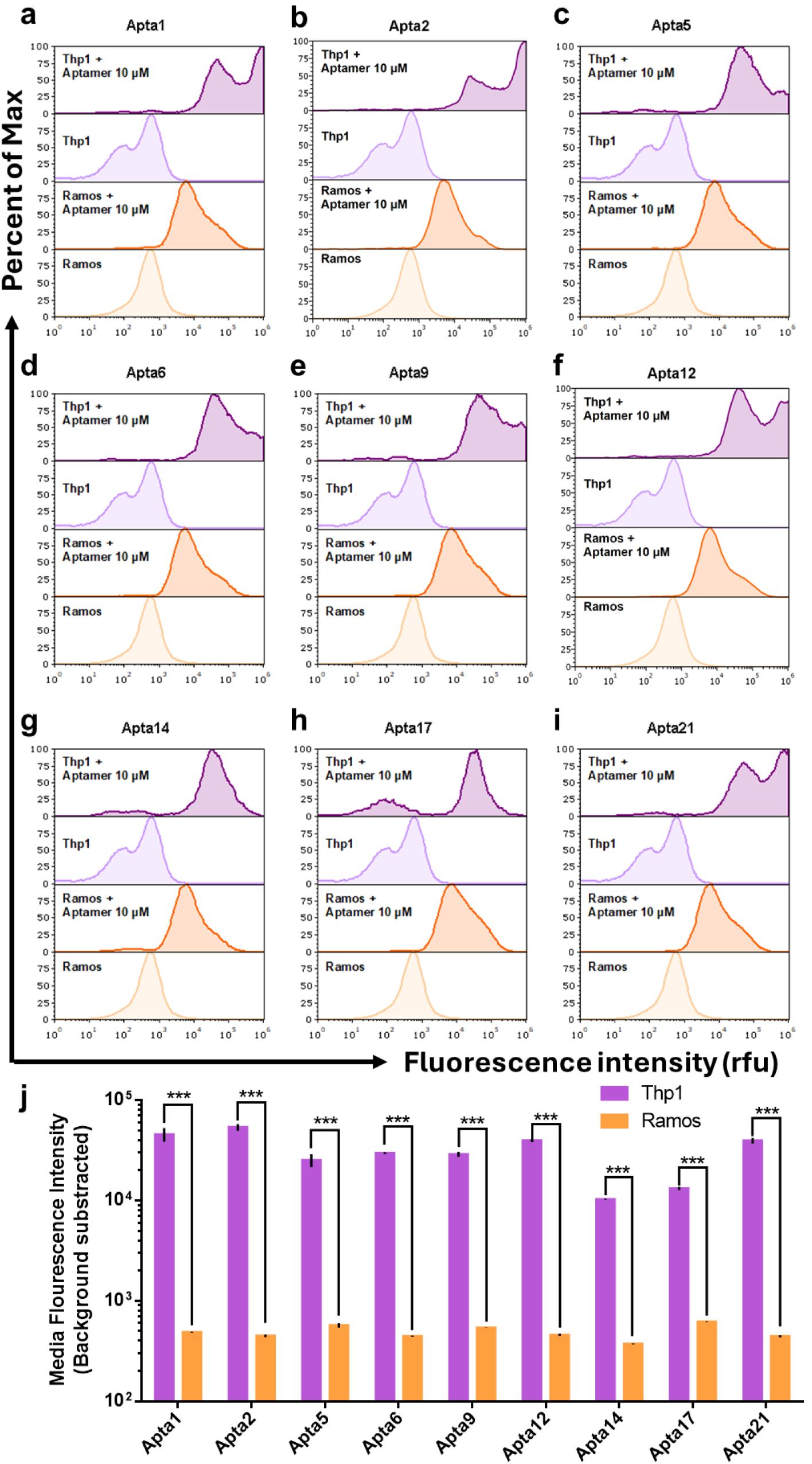
CD64 aptamers predicted by APIPredWeb demonstrate specific cellular binding. (a-i) Flow cytometry histogram profiles comparing binding of nine APIPredWeb predicted aptamers to Thp1 and Ramos cells 1 h post treatment. Untreated cells serve as background controls. X-axis represents Fluorescence in rfu and Y-axis represents Percent of Max cell population. **(j)** Quantitative analysis of background subtracted signals highlighting the specificity of the aptamers to Thp1 cells (express CD64) against Ramos cells (don’t express CD64) *** indicates a p-value less than 0.005.

## Discussion

Aptamers offer several advantages over antibodies that include low-cost chemical synthesis, improved thermal and storage stability, reduced immunogenicity, and high manufacturing reproducibility. Despite these properties, aptamer adoption in translational and clinical pipelines remains limited, largely because aptamer discovery still relies predominantly on iterative experimental enrichment workflows (e.g., SELEX-derived protocols). These processes are time-and resource-intensive, require repeated rounds of selection and characterization, and frequently demand substantial empirical optimization to achieve the affinity–specificity profiles required for downstream deployment. As a result, aptamer discovery remains a practical bottleneck that constrains throughput and delays target-to-binder timelines. Recent advances in machine learning have opened new avenues for predicting functional high-affinity peptide and small-molecule binders, supported by strong experimental validation ^10^ ^11^ ^12^. In the aptamer space, several machine learning–based platforms, including AptaNet ^4^, PPAI ^13^, and AptaTrans ^3^, have been developed to assess aptamer–protein interactions. Building upon these efforts, we previously introduced APIPred ^9^, an aptamer–protein interaction prediction pipeline. However, existing tools primarily evaluate binding potential and lack the ability to generate novel aptamer sequences for user-defined protein targets.

APIPredWeb 1.0 addresses this limitation by leveraging interaction prediction capabilities to generate novel aptamer sequences through a user-friendly interface that supports customizable aptamer libraries with defined sequence variance. Beyond aptamer generation, APIPredWeb 1.0 is being further developed towards an active-learning framework that continuously incorporates newly reported aptamers—whether generated using APIPred or conventional approaches—to iteratively improve predictive performance.

In the context of emerging pathogens, increasing antimicrobial resistance, and the growing need for targeted and personalized cancer therapies, APIPredWeb 1.0 offers a novel and scalable platform for the rapid development of aptamer-based binders targeting clinically relevant proteins.

## Methods

### 1. Sequence Library Creation and Optimization

The platform implements a sophisticated sequence library creation algorithm that balances biological relevance with computational efficiency through advanced vectorization and batch processing optimizations. The process begins with user-defined parameters for variant length and total sequence length, followed by intelligent sequence generation that incorporates constraint-based optimization.

The optimization pipeline employs several key computational enhancements: Precomputed amino acid feature vectors are calculated once per job using the PAAC algorithm, generating a 300-dimensional feature vector that is reused across all sequence predictions. Vectorized constraint pre-filtering utilizes pre-compiled regular expressions and numpy array operations to rapidly eliminate sequences violating biological constraints, achieving order-of-magnitude speedup over iterative checking methods. Batch processing architecture groups up to 500 sequences for simultaneous evaluation, minimizing Python cost and maximizing computational throughput.

The system pre-calculates valid GC count ranges for the variable region based on fixed prefix and suffix sequences, then employs generation of base compositions within valid GC ranges. Ultra-fast constraint checking implements optimized constraint verification through pre-compiled regex patterns for consecutive base detection.

XGBoost DMatrix API integration replaces standard prediction methods with optimized DMatrix data structures, providing 2-4x speedup through efficient memory layout and reduced Python-to-C++ cost. Bounded priority queue implementation uses a min-heap data structure to maintain only the top 25 results throughout processing, with O(log 25) insertion complexity replacing O(n log n) sorting operations on potentially millions of sequences, significantly reducing memory usage and computational cost.

### 2. Structural Analysis Integration

The platform incorporates ViennaRNA Package 2.0+ for comprehensive DNA secondary structure analysis with parallel processing optimizations that significantly accelerate structural computations ^14^. ViennaRNA uses dynamic programming algorithms based on the Zuker & Stiegler approach with Turner 2004 energy parameters for thermodynamic calculations.

Parallel DNA folding implementation utilizes Python’s ThreadPoolExecutor for multithreaded computation (8 threads) to perform concurrent structure predictions across multiple sequences. Batch folding architecture groups sequences into optimal batch sizes (currently 500 sequences per batch) and distributes folding tasks across available CPU cores. The parallel system includes fallback mechanisms that ensure robust operation even when parallel processing encounters errors. The MFE calculation process uses ViennaRNA’s “RNA.fold()” function through the optimized parallel interface. The parallel implementation distributes dynamic programming calculations across multiple worker threads, performs backtracking to determine secondary structures, calculates energy contributions from stacking and loop interactions, and collects both structure strings and MFE values. Performance monitoring tracks folding speeds and automatically adjusts parallelization parameters to optimize throughput.

The structural prediction pipeline integrates seamlessly with the prediction workflow, providing orthogonal assessment of aptamer viability through thermodynamic analysis. The system considers energy model components, particularly, base pair stacking as the most stabilizing element, loop penalties that provide destabilizing contributions from hairpin, bulge, and internal loops, and terminal mismatches contributing energy from unpaired bases adjacent to helices ^14^.

### 3. Statistical Analysis Framework

The platform employs a comprehensive statistical framework that evaluates performance through multiple metrics enhanced with real-time performance analytics. Interaction probability serves as the primary ranking metric derived from XGBoost predictions using the optimized DMatrix API, while thermodynamic stability from MFE values provides an independent assessment of aptamer viability. The system analyzes sequence diversity in variable regions across top candidates and monitors constraint compliance by tracking the percentage of generated sequences meeting biological constraints.

Performance benchmarking capabilities provide comprehensive analysis of optimization effectiveness, including batch processing analytics that monitor throughput improvements, processing speeds, and resource utilization. Parallel processing statistics evaluate DNA folding speedups, worker thread efficiency, and optimal parallelization parameters.

Quality assessment incorporates biological relevance through constraint enforcement, ensuring generated sequences maintain aptamer-like properties. This framework ensures that results represent not only high-scoring predictions but also biologically meaningful and diverse aptamer candidates delivered with optimal computational efficiency.

### 4. Minimum Free Energy Analysis and Interpretation

The MFE of DNA molecules is influenced by three primary factors: sequence length, nucleotide composition, particularly GC content, and nucleotide arrangement. Longer sequences tend to be more stable due to increased stacking and hydrogen bonding opportunities, while GC-rich sequences exhibit greater stability than AT-rich sequences. ViennaRNA employs the Turner 2004 energy parameters, incorporating contributions from various structural elements where base pair stacking provides the most stabilizing interactions between adjacent base pairs, while loops contribute destabilizing effects from hairpin, bulge, and internal loop formations ^14^.

The relationship between MFE and interaction probability represents a crucial aspect of result interpretation. These measures provide independent assessments of aptamer quality. MFE reflects intrinsic thermodynamic stability of the aptamer secondary structure, while interaction probability predicts binding affinity to the target protein based on aptamer sequence and structural features. No direct correlation exists between these measures, as high interaction probability does not necessarily correspond to low MFE values. Binding affinity depends on protein-specific interactions rather than general structural stability for the RNA secondary structure predicted by ViennaRNA.

### 5. Cell culture, Flow cytometry and Statistical Analysis

THP-1 and Ramos cell lines were obtained from the American Type Culture Collection (ATCC). Aptamer oligonucleotides conjugated with a 5-FAM fluorophore at the 3′ end were purchased from Integrated DNA Technologies (IDT, USA). Aptamers were resuspended to a final concentration of 100 µM in a folding buffer containing 20 mM HEPES (pH 7.2), 100 mM NaCl, and 4 mM MgCl₂. Aptamer folding was performed in the same buffer by heating samples to 95 °C for 5 min to ensure complete denaturation, followed by rapid cooling to 4 °C to promote proper secondary structure formation.

Cells were cultured in Human Plasma-Like Medium (HPLM; Thermo Fisher Scientific, A4899101) supplemented with 10% heat-inactivated fetal bovine serum (FBS; Thermo Fisher Scientific, 10100147) and 1× antibiotic–antimycotic solution (Thermo Fisher Scientific, 15240062). Cultures were maintained at 310 K in a humidified incubator with 5% CO₂. The medium was refreshed every 48 h, and cells were passaged every 3-4 days. For routine maintenance and experimental use, cells were harvested by centrifugation at 300 × g for 5 min and resuspended as required. Cell counts were determined using an automated cell counter (Countess, Invitrogen) according to the manufacturer’s instructions.

For binding assays, approximately 1 × 10^4^ cells in 100 µL of fresh medium were aliquoted into each well of a 96-well plate. Cells were incubated with 5-FAM–labeled aptamers at a final concentration of 10 µM at 310 K in the folding buffer supplemented with 10% FBS. Untreated cells served as negative controls. Following incubation, cells were washed three times by centrifugation using ice-cold flow buffer, resuspended in ice-cold flow buffer, and analyzed on a Thermo Attune NxT flow cytometer (Thermo Fisher Scientific, A24863).

Doublets were excluded during analysis, and fluorescence signals were gated on singlet populations. Mean fluorescence intensity (MFI) values were calculated for each condition. Comparative analysis of MFI values was performed using an unpaired two-tailed *t*-test, with *p* < 0.05 considered statistically significant.

## Data and Code Availability

We run the server on a virtual machine provided by the National Center for Supercomputing Applications (NCSA) department of the University of Illinois Urbana-Champaign (UIUC), ensuring reliable computational resources and accessibility for the research community.

The APIPred code is available at https://github.com/Meaw0415/APIPred. The URL for the APIPred webserver will be provided upon reasonable request to the corresponding authors. The URL will be made publicly available following acceptance of the final manuscript.

## Acknowledgement

The authors would like to thank the National Center for Supercomputing Applications (NCSA) at the University of Illinois for offering Radiant Computational Services. The study is partially supported by the Chan Zuckerberg BioHub Spoke Award.

## Notes

https://github.com/Meaw0415/APIPred

### Competing Interest Statement

The authors have declared no competing interest.

